# Cobamide-based interactions between soil bacteria can be predicted based on monoculture growth

**DOI:** 10.1101/2025.09.16.676556

**Authors:** Zoila I. Alvarez-Aponte, Hazel I. Anagu, Kenny C. Mok, Sandy T. Nguyen, Valentine V. Trotter, Michiko E. Taga

## Abstract

Interactions between microbes shape the structure and function of microbial communities. While studying interactions is key to understanding microbial communities as a whole, gaining a detailed mechanistic view is challenging due to the number of co-occurring interactions. Here, we focus on a single class of shared nutrients, the cobalamin (vitamin B_12_) family of enzyme cofactors known as cobamides, to study nutrient interactions in laboratory consortia of increasing complexity. Measuring cobamide-dependent growth of two soil *Caulobacter* isolates in monoculture revealed that each grows faster at a distinct range of cobamide concentrations. Co-culturing the two isolates demonstrated that the cobamide concentration predictably determines which isolate predominates. These results suggest that, while the two organisms appear to be functionally redundant, with identical metabolic capabilities predicted in their genomes, they may occupy distinct niches in their environment based on cobamide concentration. We further examined the ability of cobamide-producing *Mesorhizobium* and *Priestia* isolates to share cobamides and found that both support the cobamide-dependent growth of the two *Caulobacter* isolates. The *Caulobacter* isolates could coexist in tricultures with each producer following serial passaging, indicating the producers can support the growth of multiple cobamide-dependent organisms. We analyzed the metabolic capacity encoded in the genomes of the four isolates and found that cobamides are likely the main shared nutrient in our co– and tri-cultures. These results highlight the utility of cobamides as a model nutrient to characterize and predict interactions in bacterial consortia of increasing complexity.

**Importance:** Interactions between microbes shape their communities and the world around them. Because microbial communities are physically and chemically complex, predicting microbial interactions is often difficult. Using a set of four bacteria isolated from the same soil environment, we generated and tested predictions about the interactions involving a family of model nutrients related to cobalamin (vitamin B_12_). Understanding and predicting interactions can lead to greater insights into microbiome function.

## Introduction

From soils to oceans to the human gut, microbial communities shape processes crucial for human health and the planet. There is growing interest in predicting how different biotic and abiotic factors impact microbial communities (1–5). Interactions between microbes are strong determinants of community structure and function, and single microbes or single metabolites can impact the community broadly (6–10). Thus, deciphering how microbe-microbe interactions occur and are regulated is crucial for understanding whole communities. However, microbial communities encompass complex networks with many metabolites being produced and consumed at a given time, making it difficult to observe and disentangle specific interactions (11–14).

Focusing on interactions involving a single shared nutrient enables the mechanistic study of specific microbial interactions, while setting aside other interactions that could have confounding effects (15, 16). Cobamides, the cobalamin (vitamin B_12_) family of cofactors (Fig. 1), are useful as model nutrients for several reasons. First, cobamide-sharing interactions have been observed in bacterial co-cultures (15, 17–20). Moreover, cobamides are predicted to be shared widely among bacteria, based on genomic data showing an estimated 86% of sequenced bacterial species use cobamides yet only 36% have the complete biosynthesis pathway (21). Second, they are required enzyme cofactors for diverse metabolic processes that are relevant in different environments, including metabolism of specific carbon and nitrogen sources, methionine synthesis, and reductive dehalogenation (21, 22). Third, bacteria have distinct preferences for different cobamides that impact their growth and metabolism (23). These characteristics, combined with prior foundational research on cobamide biosynthesis and use, give us the ability to study cobamide-based microbial interactions. The investigation of interactions through the lens of a model nutrient can enhance our understanding of microbial competition and cooperation in laboratory consortia and in the environment.

**Figure 1.**
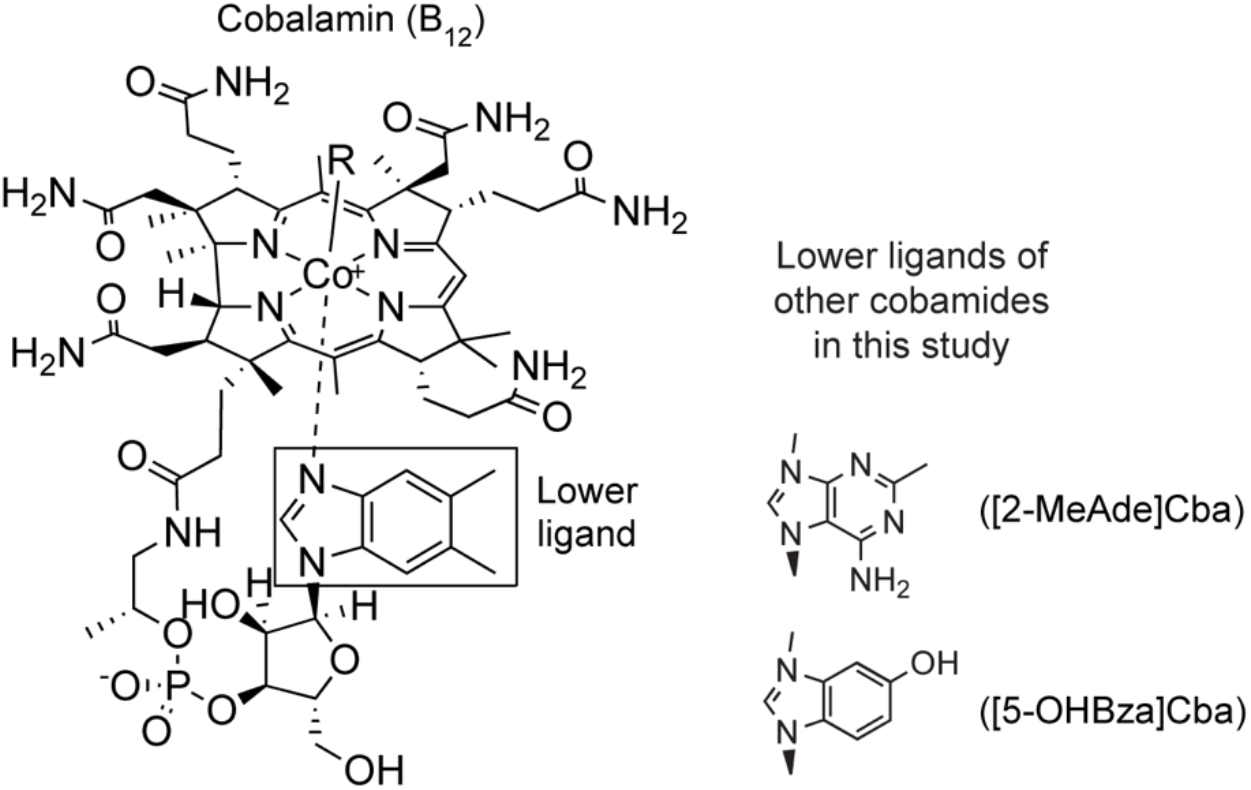
Cobamides used in this study. (Left) Structure of cobalamin (B_12_) with its lower ligand, 5,6-dimethylbenzimidazole, shown inside the rectangle. (Right) Lower ligands of 2-methyladeninylcobamide ([2-MeAde]Cba) and 5-hydroxybenzimidazolylcobamide ([5-OHBza]Cba) are shown.

We previously isolated bacteria from a grassland soil in California and characterized their cobamide metabolism (24). We classified isolates capable of producing cobamides *de novo* as “producers” and those that required cobamides for growth as “dependents” and observed that nearly all the dependents in our isolate collection preferred cobalamin over other cobamides but could use different cobamides to various extents (24). The soil isolates we evaluated required lower cobamide concentrations for growth compared to bacteria from other environments, suggesting that soil microbes have adapted to relatively low concentrations of bioavailable cobamide (23, 25). Because cobamide structure and concentration can greatly influence the growth and metabolism of bacteria (23), it is likely that competition between dependents is shaped by the identity and amount of cobamide present. Thus, we hypothesized that cobamide-dependent growth in monoculture can be used to predict population dynamics in consortia where cobamide metabolism drives interactions.

Although our soil isolate collection is taxonomically diverse, all of the cobamide producers that we isolated from soil were found to synthesize cobalamin in monoculture (24). Of this set, all contained intracellular cobalamin, but only a subset of producers had detectable amounts of cobalamin in culture supernatants. Although the mechanism of cobamide release remains unknown, cobamides have been detected in culture supernatants of many bacteria (16, 24, 26). Some producers released cobalamin at levels exceeding the requirements of the dependents, leading us to postulate that these bacteria could support the growth of dependents in co-culture (24). Further, we hypothesized that the extent to which each producer could support dependents would be predictable based on the amount of extracellular cobamide detected in monoculture.

Here, we characterized cobamide-based interactions among four bacterial isolates—two dependents with different growth profiles on cobamides and two producers that release different amounts of cobamide in monoculture (24). We grew these bacteria in co– and tri-cultures to test the hypothesis that growth and cobamide measurements in monoculture and co-culture can reliably predict community dynamics in co-culture and tri-culture, respectively. Indeed, we found that when two dependent isolates were grown in co-culture, we could predict which isolate would have a higher abundance based on their different cobamide requirements in monoculture, which may reflect the distinct niches they occupy in the environment. Further, we determined that cobamides released by producers can support the growth of dependents in co-culture.

Finally, we found that cobamide release by a producer can impact the growth of dependents in tri-cultures. The centrality of cobamide-based interactions among these soil bacteria is reinforced by a comparison of their genome sequences. Overall, these insights contribute to a better comprehension of pairwise bacterial interactions and enhance our understanding of how cobamides may impact dynamics in natural communities.

## Results

### Dependents have advantages at distinct cobamide concentrations

We hypothesized that the dominant isolate in co-cultures of two dependent bacteria could be predicted based on their relative cobamide-dependent growth rates in monoculture (23). We tested this hypothesis on a pair of cobamide-dependent soil isolates, *Ca*19 and *Ca*24A, initially selected because they grew reproducibly with no aggregation in liquid culture (Fig. S1A, B). Through the genome sequencing effort described in a later section, we found that the isolates belong to two different species of *Caulobacter*, as reflected by a whole genome average nucleotide identity (ANI) of 89% (27). The species were not identified by the Genome Taxonomy Database Toolkit (GTDB-Tk) (28, 29), which suggests that they could be novel species.

We measured the maximum growth rates of *Ca*19 and *Ca*24A in media containing a range of concentrations of [5-OHBza]Cba, cobalamin, and [2-MeAde]Cba (Fig. 1) to determine how these cobamides influence growth in monoculture (Fig. 2A, D, G). The dose response curves for the two isolates grown on [5-OHBza]Cba and cobalamin are distinct, as demonstrated by statistically significant differences in one or more curve parameters (Fig. S2). *Ca*19 has a higher growth rate than *Ca*24A at low concentrations of [5-OHBza]Cba, whereas *Ca*24A grows faster at high concentrations (Fig. 2A). We observed a similar trend on cobalamin, although the differences were less pronounced (Fig. 2D). The cobamide concentration required to reach the half-maximal growth rate (EC_50_) was significantly lower for *Ca*19 on these two cobamides (p < 0.05 for cobalamin, p < 0.001 for [5-OHBza]Cba) (Fig. S2A), suggesting this isolate may be adapted to lower cobamide concentrations. In contrast, the two isolates grow at similar rates at all concentrations of [2-MeAde]Cba, and no statistically significant differences were observed for the dose response curve parameters when isolates were grown with this cobamide. (Fig. 2G, S2).

**Figure 2.**
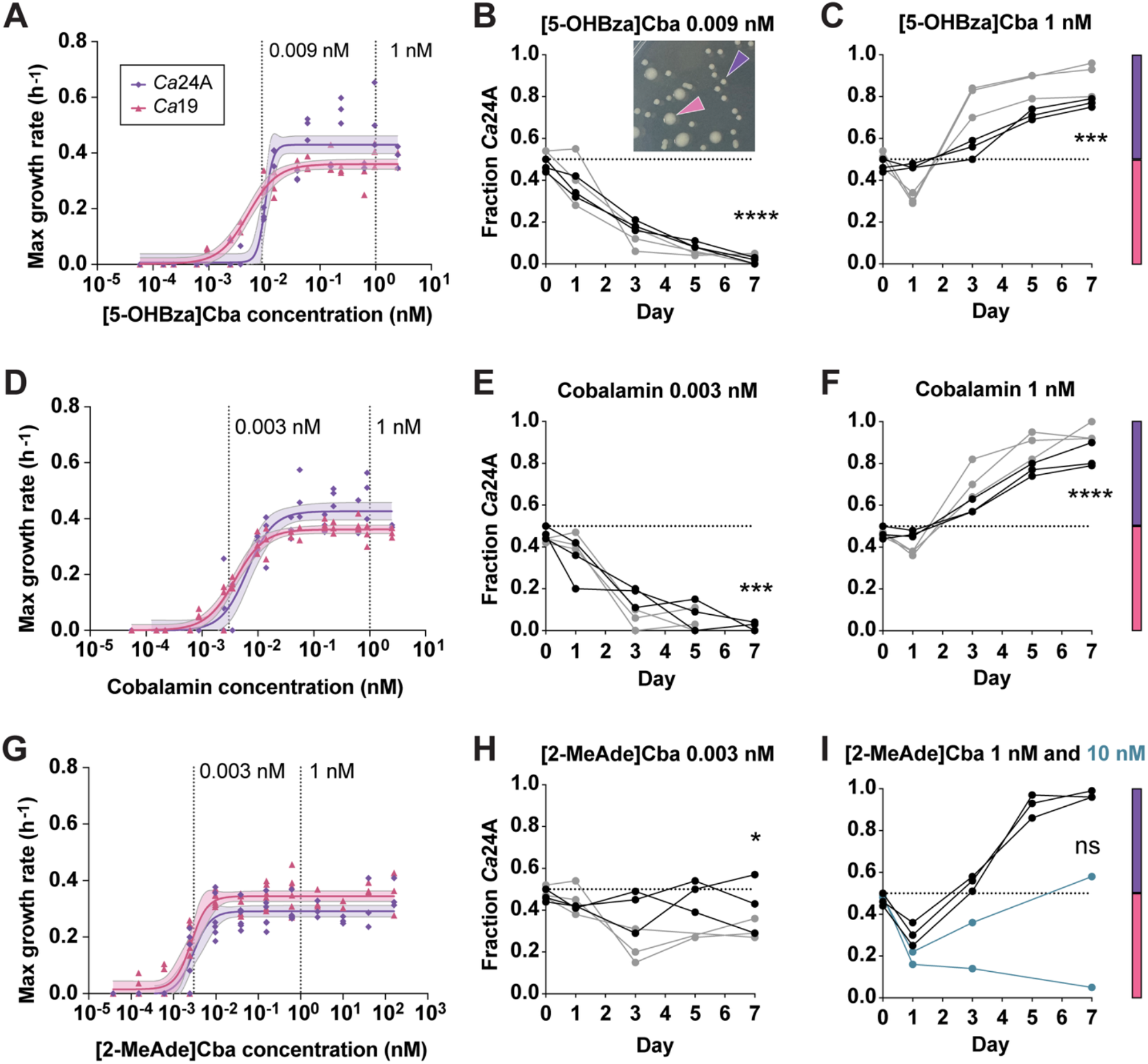
Dependent isolate growth rates in monoculture predict final dependent abundances at different cobamide concentrations in co-culture. The maximum growth rate of dependent monocultures grown in a range of concentrations of (A) [5-OHBza]Cba, (D) cobalamin, and (G) [2-MeAde]Cba is shown. Individual data points are shown for each biological replicate (n=6), with three no-growth outliers removed. Curves represent a four-parameter least squares fit and 95% confidence interval, and dotted vertical lines denote the concentrations selected for further experiments. (B, C, E, F, H, I) Graphs show the relative abundance of *Ca*24A in co-culture with *Ca*19, passaged daily for seven days, based on the number of colony forming units (CFU/ml) of each isolate in samples taken at the indicated time points. (B, C) [5-OHBza]Cba, (E, F) cobalamin, and (H, I) [2-MeAde]Cba were provided at (B, E, H) low and (C, F, I) high concentrations, as indicated. Lines show data for independent (n=5 or n=6) biological replicates from two independent experiments (black lines and gray lines) performed under the same conditions, except that a higher concentration of [2-MeAde]Cba (I) was used in the second experiment for the high-concentration of this cobamide (blue lines). The dotted horizontal line indicates 50% abundance, and the color bars indicate the dominating isolate at the corresponding region of the graphs (purple for *Ca*24A and pink for *Ca*19). Asterisks denote the result of a one-sample t and Wilcoxon test used to evaluate if there was a statistically significant difference between the mean value of the last timepoint and a hypothetical value of 0.5. One asterisk represents p<0.05, two asterisks p<0.01, three asterisks p<0.001, and four asterisks p<0.0001, while ‘ns’ indicates non-significant (p>0.05). The inset in B shows different colony morphologies of the two dependents grown in co-culture: *Ca*24A (small colonies, purple arrowhead) and *Ca*19 (large colonies, pink arrowhead) after 72 hours of growth on R2A agar containing cobalamin. Statistical comparisons for the curves shown in A, D, and G can be found in Figure S2.

We expected that, given that the cobamide concentration can determine growth rate in monoculture, the isolate with the higher growth rate at a given cobamide concentration should outcompete the other in co-culture. We tested this hypothesis by establishing co-cultures at different cobamide concentrations and differentiating between the two isolates based on their distinct colony morphologies (Fig. 2B, inset). Based on their relative growth rates in different concentrations of [5-OHBza]Cba and cobalamin, we predicted that *Ca*19 would outcompete *Ca*24A when co-cultured at concentrations below 0.012 nM [5-OHBza]Cba or 0.010 nM cobalamin (the approximate points of intersection between the two curves), whereas *Ca*24A should outcompete *Ca*19 at higher cobamide concentrations (Fig. 2A, D). While growth rate differences were minor in [2-MeAde]Cba, there was a small growth advantage for *Ca*19 (Fig. 2G). Therefore, we predicted that *Ca*19 would outcompete *Ca*24A at all concentrations of [2-MeAde]Cba. We co-cultured *Ca*19 and *Ca*24A in media containing a low (0.003 or 0.009 nM) or high (1 or 10 nM) concentration of each cobamide and passaged every 24 hours. The viable cell number of each isolate was determined for the inocula and co-cultures on days 1, 3, 5, and 7 (Fig. 2B, C, E, F, H, I).

Predictions of the final relative abundances of each dependent in co-culture were accurate in the low and high concentrations of [5-OHBza]Cba and cobalamin, with *Ca*19 becoming dominant at low cobamide concentrations and *Ca*24A becoming dominant at high concentrations (Fig. 2B, C, E, F). These results were consistent with our hypothesis that the dominant isolate could be predicted from maximum growth rates in monoculture, which suggests that differences in growth rate in different cobamide concentrations are the main factor driving dynamics in these co-cultures. Overall, our data suggest that *Ca*19 is adapted to lower concentrations of [5-OHBza]Cba and cobalamin and *Ca*24A is adapted to higher concentrations, and these adaptations determine their dynamics in co-culture.

In media containing [2-MeAde]Cba, final dependent isolate abundances were not accurately predicted from monoculture growth rates at the low or high concentrations (Fig. 2H, I). The isolates remained near equal proportions in the low concentration, while *Ca*24A was more abundant in the high concentration condition. Because differences in growth rates were small, we decided to evaluate a higher concentration of [2-MeAde]Cba. When the experiment was repeated with 10 nM [2-MeAde]Cba, the results differed from the 1 nM condition and the two replicates diverged (Fig. 2I, gray lines represent the 10 nM condition). Thus, due to insufficient differences in growth rate and EC_50_, predictions based on monoculture growth were not reliable for this cobamide.

### Producer supernatants support growth of dependents in monoculture

Next, we hypothesized that each dependent isolate could be supported by a producer that releases cobamides into the culture media, and that the amount of cobamide the producer provides would impact the relative abundances of the two dependents in tri-culture. We selected two isolates from the same collection—*Me*12, a strain of *Mesorhizobium opportunistum*, and *Pr*24B, a strain of *Priestia megaterium*—that were previously found to produce and release different amounts of cobalamin (24). *Pr*24B was originally co-isolated with the dependent *Ca*24A, suggesting these bacteria may interact in their natural soil environment, including via cobamide sharing. These producers synthesize and release cobalamin at concentrations that exceed the minimal requirements of both dependents in monoculture (24).

To evaluate cobalamin release in producer cultures, we prepared cell-free supernatants from producers during exponential and stationary phases of growth (Fig. S1C, D, E) and measured growth of the two dependent isolates in medium containing these supernatants. In control cultures lacking cobalamin, growth of both dependents was minimal (Fig. 3, NA). *Me*12 exponential phase supernatants caused a slight increase in dependent growth, which was only statistically significant for *Ca*19 (Fig. 3A, B). The *Me*12 stationary phase supernatants, which were diluted to avoid saturation of dependent growth, led to a significant increase in growth for *Ca*19 to a level that was not statistically distinguishable from growth with saturating cobalamin, despite the supernatant being diluted (Fig. 3B). In contrast, *Me*12 stationary phase supernatants led to a nonsignificant increase in growth of *Ca*24A, but when adjusted for the dilution factor, would represent a considerable growth increase (Fig. 3A).

**Figure 3.**
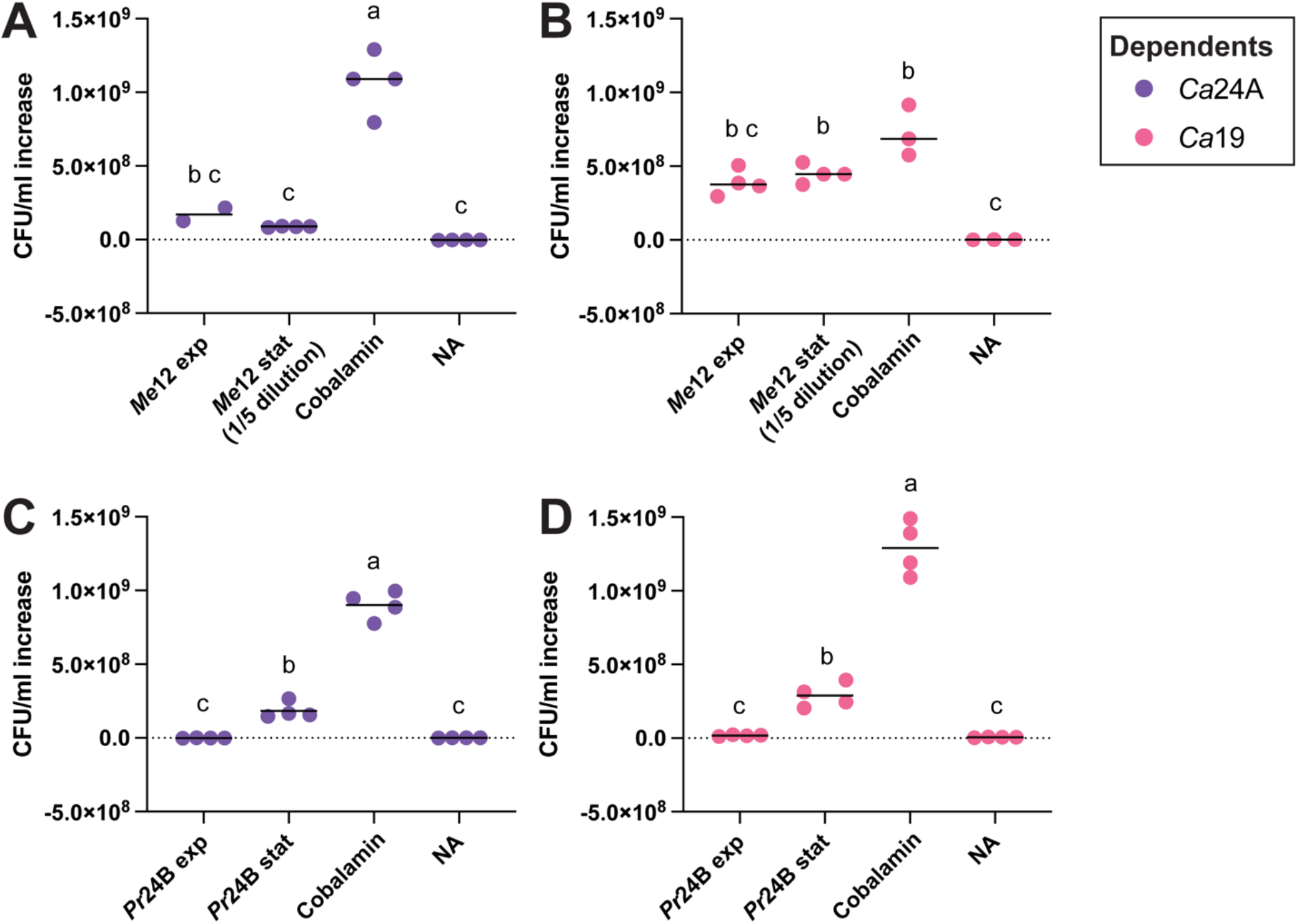
Growth of cobamide dependent isolates in medium amended with producer supernatants. Graphs show CFU/ml increase (final CFU/ml – initial CFU/ml) of dependent cultures supplemented with exponential (exp) or stationary (stat) phase supernatants from (A, B) *Me*12 or (C, D) *Pr*24B. 1 nM cobalamin addition and no addition (NA) controls are shown for each experiment. All cultures were grown for 24 hours. Points represent individual biological replicates (n=4) except for cobalamin samples in B, where n = 3. Horizontal lines show the mean CFU/ml increase for each isolate. Datapoints were compared via ordinary one-way ANOVA followed by Šídák’s multiple comparisons test, with a single pooled variance. Groups with different letters were significantly different from one another (p < 0.05), whereas groups sharing a common letter were not significantly different.

*Pr*24B supernatants prepared from cultures in stationary phase, but not exponential phase, significantly enhanced growth of both dependents (Fig. 3C, D). These results suggest that cobamide release occurs differently in the two producers. We ruled out the possibility that the producers instead provide methionine, a metabolite that commonly requires cobamides for production (21), by showing that methionine addition did not rescue growth of *Ca*24A and *Ca*19 (Fig. S3). Further, we previously showed that cobamide producer cell lysates do not contain sufficient methionine to promote growth of an *E. coli* methionine auxotroph (24).

### Producers can support dependents by providing a cobamide in co-culture

Having found that producer supernatants can fulfill the cobamide requirements of the dependents, we next tested whether producers could support dependent growth when cultured together. We hypothesized that in co-culture, release of cobalamin by the producer would directly enable dependent growth, which in turn could allow the dependents to compete more effectively for the other resources in the medium. Thus, we investigated how each isolate grew in producer-dependent co-cultures. We found that both dependent isolates grew significantly more when co-cultured with *Me*12 than in monoculture without cobalamin supplementation, with *Ca*24A reaching abundances similar to those in monocultures supplemented with a saturating concentration of cobalamin (Fig. 4A). *Me*12 provided cobalamin despite being at a significant competitive disadvantage in co-culture, reaching a density around 4-fold lower than when grown alone (Fig. 4A). When co-cultured with *Pr*24B, both dependents grew to higher levels than in monocultures with no cobalamin, but this difference was not significant (p > 0.05); their final densities were over 10-fold lower than when grown in monoculture with saturating cobalamin (Fig. 4B). Furthermore, unlike *Me*12, *Pr*24B grew to a similar level in co-culture and in monoculture, perhaps because it experienced less competition with the dependents due to the restricted cobamide availability (Fig. 4B).

**Figure 4.**
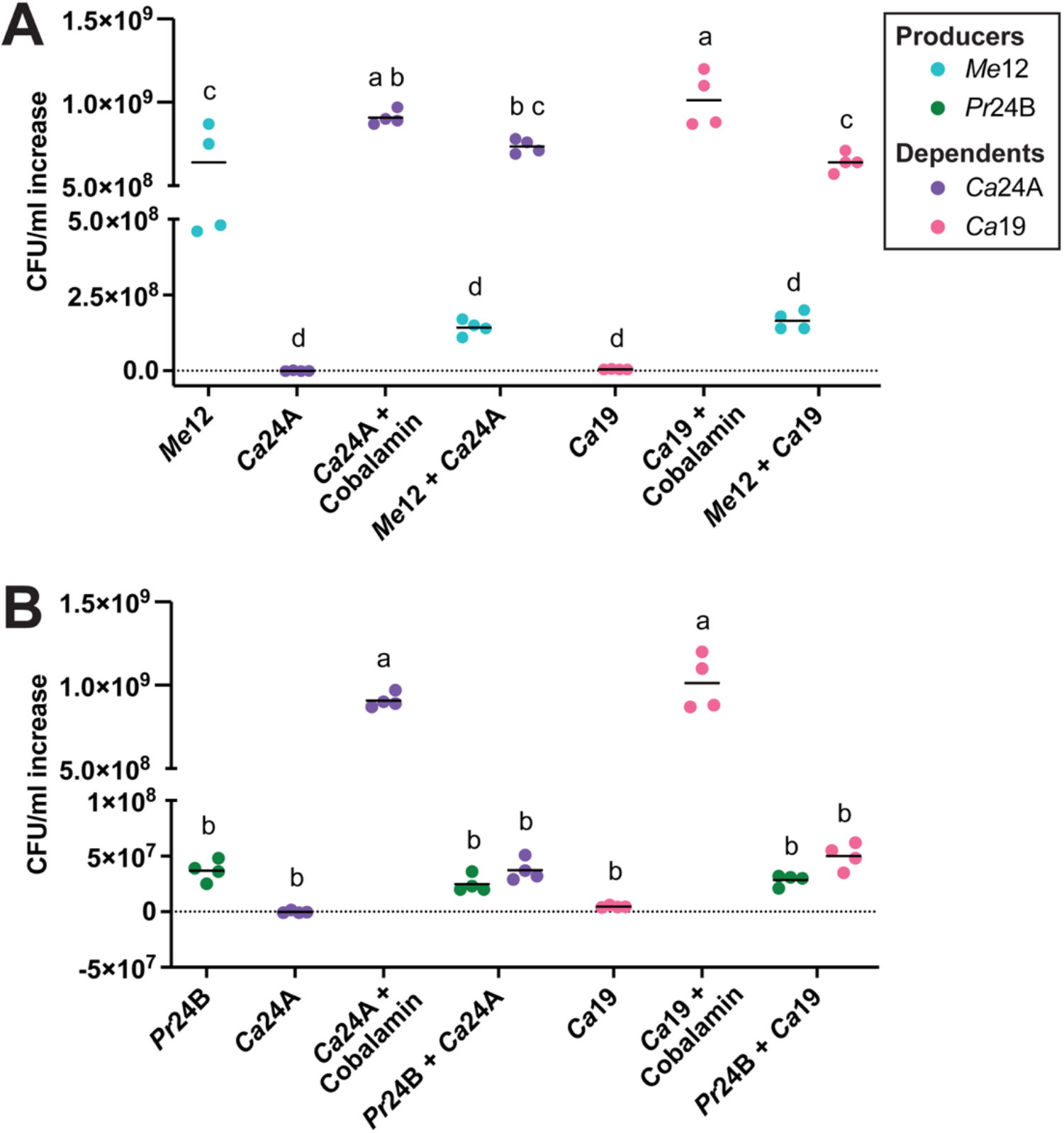
Growth of cobamide dependent isolates in co-culture with a producer. The increase in CFU/ml (final CFU/ml – initial CFU/ml) is shown for monoculture controls in medium lacking cobalamin or containing 1 nM cobalamin and co-cultures of each dependent with (A) *Me*12 and (B) *Pr*24B. For clarity, dependent monoculture data are shown in both panels. All cultures were grown for 48 hours. Points represent individual biological replicates (n=4) and horizontal lines show the mean CFU/ml increase for each isolate. Isolate growth was compared across conditions via an ordinary two-way ANOVA with main effects only, followed by Šídák’s multiple comparisons test, with a single pooled variance. Groups with different letters were significantly different from one another (p < 0.05), whereas groups sharing a common letter were not significantly different.

Upon comparing the number of dependent cells supported per producer cell, we found that dependents generally benefited more from being supplemented with supernatants than from growing in co-culture (Table 1). The calculated ratios show that though *Pr*24B supernatants supported fewer total dependent cells than *Me*12 supernatants (Fig. 4), they supported significantly higher numbers of dependent cells per producer cell than *Me*12 (Table 1). The absolute cell numbers of *Pr*24B in both monoculture and co-culture were much lower than *Me*12, possibly leading to lower total accumulated cobamide (Fig. S1E). The difference in producer cell number can partially explain why the dependents grew less in *Pr*24B supernatants and in co-culture with *Pr*24B.

**Table 1.** Number of dependent cells supported per producer cell when grown with supernatants from the exponential phase (supernatant exp) or stationary phase (supernatant stat) and in co-culture.

| Producer | Condition | Dependent cells per producer cell<sup>a</sup> |  |
| --- | --- | --- | --- |
|  |  | *Ca19* | *Ca24A* |
| ***Pr24B*** | supernatant exp | 41 ± 24 | < 0<sup>b</sup> |
|  | supernatant stat | 130 ± 57 | 81 ± 17 |
|  | co-culture | 1.7 ± 0.64 | 1.5 ± 0.17 |
| ***Me12*** | supernatant exp | 32 ± 4.2 | 14 ± 7.4 |
|  | supernatant stat | 20 ± 4.0 | 4.0 ± 0.30 |
|  | co-culture | 3.9 ± 0.88 | 5.3 ± 1.0 |

<sup>a</sup>Ratios were analyzed separately for each dependent isolate using aligned rank transform ANOVA, followed by Benjamini-Hochberg-adjusted planned comparisons. For *Ca19*, significant differences between producers (*Me12* versus *Pr24B*) were observed in co-cultures ( $p < 0.005$ ), and in the stationary phase supernatant condition ( $p < 0.001$ ), but not in the exponential phase supernatant condition ( $p > 0.05$ ). For *Ca24A*, significant differences between producers were observed in co-cultures ( $p < 0.005$ ) as well as exponential and stationary phase supernatants ( $p < 0.0001$ in both cases).
<sup>b</sup>Less than 0 ( $< 0$ ) means that the dependent CFU/ml count was lower at the end than at the beginning, i.e. no dependent growth was detected.

### Cobamide providing and competition occur in three-species consortia

Having found that each producer could support both dependents, we sought to determine how producers influence the growth of the two dependents when grown in small consortia. Thus, we established tri-cultures containing the two dependents with each producer in a cobamide-free medium with daily passaging into fresh medium. We hypothesized that the amount of cobalamin released by each producer would determine which dependent was more abundant at the end of the 7-day tri-culture. On days 1, 3, 5, and 7, the cultures were plated to determine the abundances of each isolate based on their unique colony morphologies (Fig. 5A, B).

**Figure 5.**
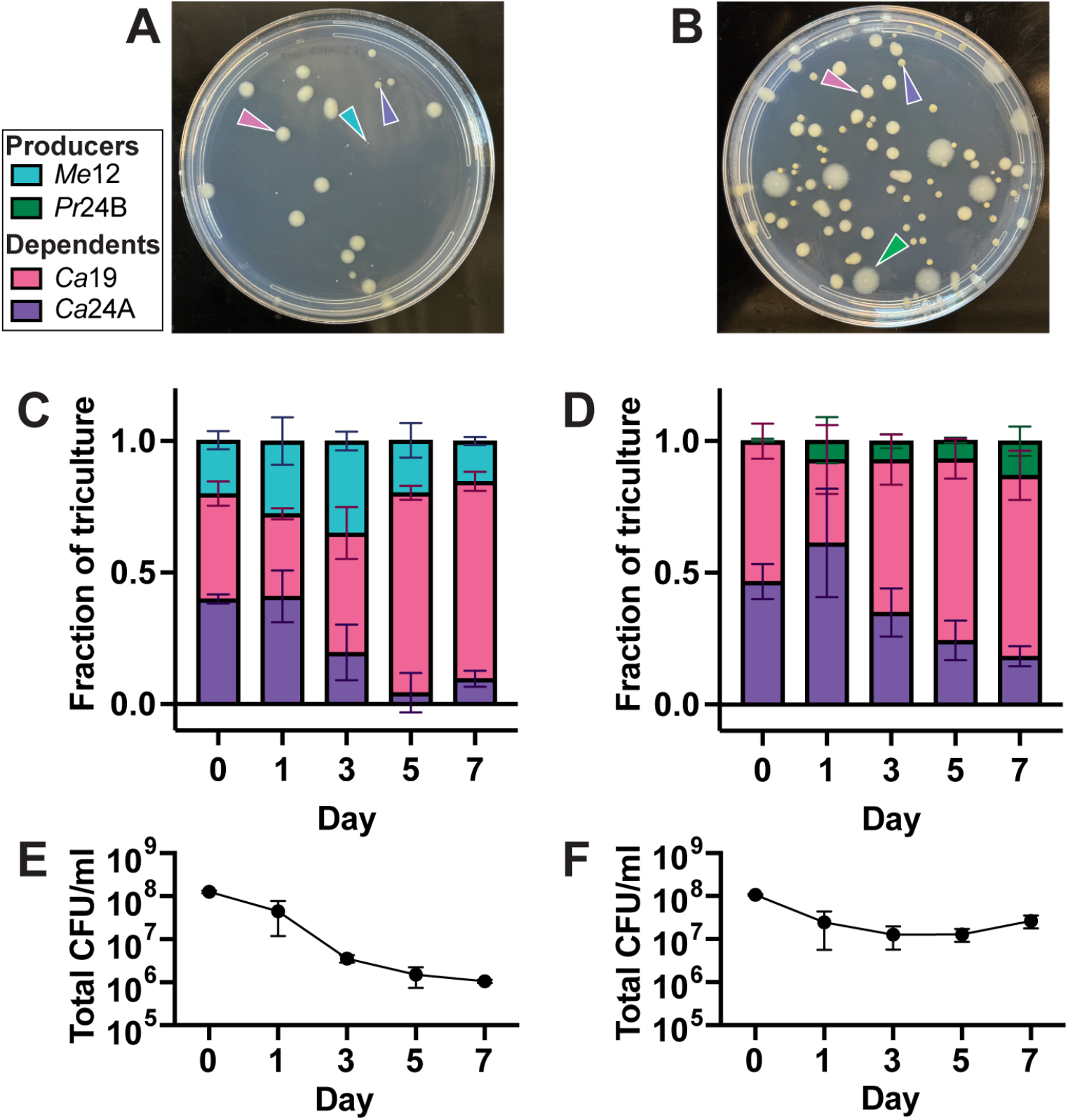
Growth of *Ca*19 and *Ca*24A in tri-culture with each producer. (A, B) Colony morphologies of both dependent isolates with (A) *Me*12 and (B) *Pr*24B after 96 and 72 hours of growth on solid medium, respectively. Arrow colors correspond to the color for each isolate in the legend. (C, D) Relative abundances of producer and both dependents in tri-culture with (C) *Me*12 and (D) *Pr*24B are shown. Bars represent the mean of three biological replicates and error bars represent standard deviation. (E, F) Total CFU/ml of the (E) *Me*12 and (F) *Pr*24B tri-cultures, respectively. Points show the mean and error bars show standard deviation.

In both tri-cultures, all three isolates persisted over the seven passages, and the producer had the lowest abundance at most timepoints (Fig. 5C, D). While *Me*12 had similar relative abundances at the beginning and end of the experiment, *Pr*24B was barely detectable at the initial timepoint and increased to over 10% by day 7. *Ca*19 was the more abundant of the two dependents by day 3 in both tri-cultures, and *Ca*19 abundance further increased at later timepoints relative to *Ca*24A (Fig. 5C, D). While the three isolates coexisted in the *Pr*24B tri-culture with total CFU/ml remaining relatively constant over time, total cell numbers in the *Me*12 tri-culture decreased with each passage (Fig. 5E, F). This likely happened because this producer has the lowest growth rate of the four isolates, requiring approximately 28 hours to reach stationary phase (Fig. S1C), and tri-culture passages were done every 24 hours, likely preventing it from reaching higher cell densities between passages.

We performed an *E. coli*-based bioassay to determine the total cobalamin concentration in our tri-cultures (Table S1) (30). We relied on whole culture cobalamin measurements because supernatant concentrations were below the detection limit of our bioassay. In the *Me*12 mono– and tri-cultures, cobalamin levels were below the bioassay detection limit (0.27 nM) at all timepoints, consistent with *Me*12 lacking sufficient time to grow and produce cobamides and decreasing total cell numbers (Fig. 5E). The *Pr*24B tri-cultures had a stable cobalamin concentration of approximately 2 nM over multiple passages, which matched the concentration sustained in *Pr*24B monocultures passaged in the same medium (Table S1).

### Genome analysis and experimental validation suggest cobamides are the main shared nutrient in these consortia

To investigate the possibility that additional metabolic requirements and capabilities could impact growth of the isolates in co– and tri-cultures, we sequenced and analyzed the genomes of the four isolates to explore whether they could produce other nutrients that are commonly shared, such as amino acids and vitamins (Fig. 6). We also considered the possibility that certain isolates could degrade carbon sources that others could not, making byproducts available, so we examined the catabolism pathways for the four carbon sources present in our medium. Further, we conducted an in-depth analysis of cobamide-related genes to examine the genetic basis of the cobamide production and dependence phenotypes in our isolates. Our genomic analysis is summarized in Table S2.

**Figure 6.**
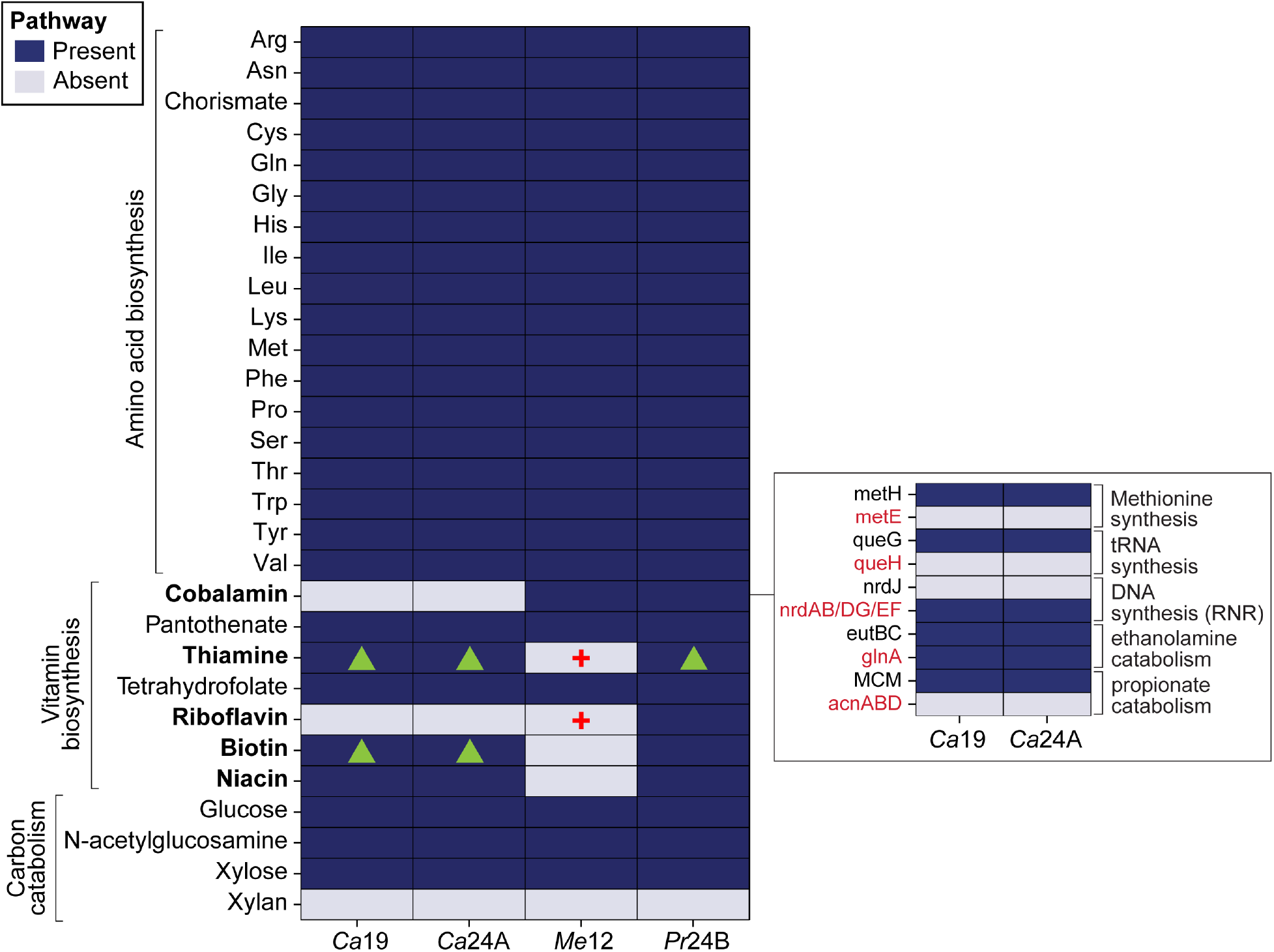
Amino acid, vitamin, and carbon catabolism pathways present or absent in the genome of each isolate. Amino acid biosynthesis and carbon catabolism were annotated using GapMind (57) and refined manually. Vitamin biosynthesis pathway presence and absence were determined by using BlastKOALA (56) to obtain KEGG Orthology (KO) annotations for each genome and cross-referencing with previously published lists of KO annotations for vitamin biosynthesis (25, 31, 32). Vitamin pathway annotations were manually reannotated with more lenient search parameters based on enzyme commission (EC) numbers. Vitamin names in bold lettering denote cases in which we experimentally evaluated bacterial growth in the presence and absence of that vitamin (experimental data can be found in Figure S4). In cases where experimental results did not align with genomic predictions, red crosses are shown. Green triangles denote vitamins for which genomic predictions based on different annotation methods were inconsistent. In the box, we show the presence or absence of cobalamin-dependent genes (black text) and their cobalamin-independent alternatives (red text) in the dependent isolate genomes. This includes genes that were relevant in our growth conditions; the full cobamide genome analysis can be found in Figure S5. RNR, ribonucleotide reductase.

Analysis of amino acid, cofactor, and carbon substrate biosynthesis, dependence, and catabolism in our genomes suggests that cobalamin is the main shared nutrient in our experimental conditions. All isolates were predicted to be capable of synthesizing all 20 proteinogenic amino acids (Fig. 6). The four isolates also have pathways for the catabolism of three of the four carbon sources in our medium (24); they share the capability to degrade glucose, N-acetylglucosamine, and xylose, but lack xylan catabolism genes (Fig. 6).

*Ca*19 and *Ca*24A are both predicted to synthesize all evaluated vitamins except cobalamin and riboflavin, while *Me*12 was predicted to lack complete pathways for thiamine, riboflavin, biotin, and niacin biosynthesis, and *Pr*24B is predicted to have complete biosynthesis pathways for all the evaluated vitamins (Fig. 6). While this may result in *Pr*24B providing additional nutrients or having a competitive advantage in co-culture, all of these vitamins were present in the culture medium and likely did not largely impact the outcome of co– and tri-cultures. Because riboflavin was present in the culture medium, cobalamin is likely to be the main shared nutrient in our co-cultures and tri-cultures.

We used two annotation methods to predict vitamin auxotrophy, and results differed for thiamine and biotin (31, 32). Therefore, to experimentally evaluate the isolates’ vitamin requirements, we passaged each isolate in biotin-, niacin-, thiamin-, or riboflavin-deplete medium, expecting that isolates not predicted to synthesize a particular vitamin would fail to grow in that vitamin’s absence (Fig. S4). Our results showed that the three isolates with incongruous predictions for thiamine and biotin auxotrophy could grow in their absence (Fig. 6 green triangles, Fig. S4). Further, analysis of the *Me*12 genome predicted it to be auxotrophic for thiamine and riboflavin, but our experimental results showed it could grow without supplementation of either vitamin (Fig. 6 red crosses, Fig. S4). This result emphasizes the importance of validating genome-based predictions.

We further characterized the cobalamin biosynthesis and dependence genes in the four genomes to understand production and dependence of this key nutrient in our isolates. This analysis confirmed that *Me*12 and *Pr*24B are predicted to produce cobalamin while *Ca*24A and *Ca*19 lack most of the biosynthesis pathway, aligning with the phenotypic observations reported previously (Fig. 6, S5A, B) (21, 24). Further, *Ca*19 and *Ca*24A encode the same set of cobamide biosynthesis genes, dependent genes and independent alternatives (Fig. 6, expanded in Fig. S5C). Methionine was not added to our medium, as its biosynthesis is often cobamide-dependent (24). In fact, only *Pr*24B encodes the cobamide-independent methionine synthase, while *Ca*19, *Ca*24A, and *Me*12 are predicted to require cobamides for methionine production (Fig. S5C).

*Me*12 and *Pr*24B both encode *bluB*, the gene required for aerobic biosynthesis of the lower ligand of cobalamin, 5,6-dimethylbenzimidazole (Fig. 1), in accordance with our previous observation that both synthesize cobalamin (Fig. S5A-B) (24, 33, 34). *Pr*24B encodes several anaerobic corrin ring biosynthesis enzymes, suggesting it may produce cobamides via the anaerobic pathway despite being cultured aerobically in our study. The two dependent genomes lack corrin ring and lower ligand biosynthesis genes, while they contain genes for tetrapyrrole precursor biosynthesis, which are not specific to cobamide biosynthesis (Fig. S5B). However, both dependent genomes contain adenosylation, nucleotide loop assembly, and lower ligand attachment genes, suggesting they can salvage late cobamide precursors (21, 35, 36). The latter observation aligns with previous experimental results that showed both *Ca*24A and *Ca*19 can grow when provided with cobinamide, an incomplete cobamide lacking the nucleotide loop and lower ligand (24). Taken together, the analysis of the genomes of the dependents and producers in this study aligns with our experimental results demonstrating cobamide-based interactions.

## Discussion

Cooperation and competition between pairs of bacteria influence higher order interactions and shape microbial community structure (37–40). This is reinforced by the recent suggestion that microbiome health could be predicted based on microbial interactions alone (41). Achieving a mechanistic understanding of microbial communities is crucial for developing strategies for guiding communities to desired states, yet challenging because many factors contribute to microbial community interdependence (14). Here, by focusing on interactions involving cobamides, we have shown that growth measurements in monoculture can inform predictions of community dynamics in two– and three-member laboratory communities. Our finding that two *Caulobacter* species with identical metabolic gene complements can have distinct cobamide concentration optima suggests that small differences in functionally redundant taxa can contribute to community resilience. Further, the ability of two cobamide-dependent bacteria to coexist with a cobamide producer in triculture provides a platform for generating higher-order synthetic consortia.

It is curious that although the two cobamide-dependent *Caulobacter* species completely overlap in the amino acid, vitamin biosynthesis and cobamide-dependent genes they encode (Fig. S5), they have distinct cobamide concentration optima. Like adaptations for use of different cobamides, adaptations for specific cobamide concentration ranges could occur due to differences in regulation or efficiency of cobamide transporters, riboswitches, adenosylation enzymes or cobamide-dependent enzymes (16, 42). The distinct preferences of the two isolates for different cobamide concentrations illustrates how related taxa with similar metabolic capabilities can occupy distinct niches. Niche specialization based on cobamide concentration may enable coexistence of closely related bacteria in the soil environment, where nutrient availability varies across microenvironments and seasons (23, 25, 43, 44). The observation that bacteria have preferences for specific cobamide concentrations, in addition to preferred structures, adds a layer of complexity to predictions of cobamide metabolism. Because these preferences cannot be predicted from genome data, this finding emphasizes the importance of combined genomic and experimental approaches for comprehensively studying microbial nutrient-based interactions.

Our experimental evaluation of cobamide-based interactions in co– and tri-cultures revealed that cobamides, and in turn cobamide producers, can influence bacterial community dynamics. The cobamide sharing interactions we observed in co-cultures were preserved in tri-cultures, suggesting that cobamide sharing is maintained as community complexity increases (Fig. 4, 5). Predicting isolate abundances in tri-cultures based on cobamide concentrations was limited by the sensitivity of our *E. coli*-based bioassay. In the tri-culture containing *Pr*24B, total cobamide concentrations were consistently high, but the dominant isolate was *Ca*19, which grows faster at low cobamide concentrations. This could be possible if (1) *Pr*24B only released a small fraction of the cobalamin it produced, (2) cobalamin release primarily occurred at late stages of growth—as we observed (Fig. 3C, D)—and *Ca*19 benefited from low cobalamin concentrations in early stages, or (3) other interactions emerged that benefited *Ca*19 over *Ca*24A, such as the presence of other nutrients or growth inhibition of *Ca*24A. Interactions between *Ca*24A and *Pr*24B were of particular interest because they were isolated in the same culture, suggesting they could have a pre-established metabolic partnership. They were indeed able to grow together in co-culture when *Pr*24B provided cobalamin, though *Ca*24A did not have an advantage over *Ca*19 when cultured with *Pr*24B.

How producers share nutrients, whether via secretion, lysis, or contact-dependent mechanisms, remains an open question (45, 46). Cell death, particularly following phage infection, has been shown to contribute to nutrient sharing in *in vitro* systems (47, 48). It is also possible that cobamides are secreted via bidirectional transporters (49), or that they leak out of cells. It is not yet known which mechanism *Me*12 and *Pr*24B use to release the cobalamin they produce. However, we note that, despite the three isolates being inoculated at equal cell densities—estimated based on optical density—to establish the tri-cultures, viable *Pr*24B cells were barely detectable at the initial time point, suggesting most cells were dead (Fig. 5D). Thus, we speculate that cell death contributes to cobalamin release by *Pr*24B.

Our combined genomic and experimental analysis of the two *Caulobacter* species indicates that the only cobamide-dependent enzymes relevant to our experimental conditions are methionine synthase (MetH) and the tRNA modification enzyme QueG. Because methionine supplementation did not rescue growth, albeit at a single concentration (Fig. S3), we hypothesize that the epoxyqueuosine reductase QueG could be important for growth in these *Caulobacter* isolates, unlike in *E. coli,* where a *queG* mutant showed no phenotype (50). Our results also suggest *Me*12 may engage in reciprocal metabolic interactions because it provides cobalamin to other microbes but requires an external supply of biotin and niacin (Fig. 6, S4).

While cobamides were the main drivers of interactions in this simplified laboratory system, the present study is limited to four isolates, and we are likely missing important environmental signals and interactions the organisms experience in their native soil. Nonetheless, we were able to predict the outcome of dependent-dependent and producer-dependent interactions based on their growth in monoculture. This represents a significant advance in the applicability of cobamides as model shared nutrients (16). Future studies should test the predictability of cobamide interactions between additional pairs of bacteria, evaluate additional nutrient interactions, and focus on systems that more closely mimic the soil environment. Further, elucidation of the mechanisms of cobamide release may enable genome-based predictions of producer-dependent interactions. This research adds to a body of literature that has shown that predictions about nutrient sharing are possible from a combined analysis of metagenomes, isolate genomes, and phenotypic data (18, 31, 32, 38), and that nutrient availability impacts microbial competition and coexistence (51).

## Materials and Methods

### Isolate characterization and selection

The isolates used in this study were previously published as 5OH_024A (*Ca*24A), CRE_019 (*Ca*19), NOC_012 (*Me*12), and 5OH_024B (*Pr*24B) (24). The four isolates had distinct colony morphologies, making it possible to determine their abundances in co– and tri-cultures based on colony numbers.

For all experiments, bacteria were cultured at 28°C with aeration in liquid VL60 medium containing four carbon sources (glucose, N-acetylglucosamine, xylose, and xylan) and all amino acids except methionine, as described previously (24), or at 28°C on Reasoner’s 2 (R2) agar supplemented with 1 nM cobalamin. Dependents were pre-cultured by inoculating from a colony and growing to saturation in liquid medium containing 1 nM cyanocobalamin and transferring to medium lacking cobamide at a 1:100 dilution after 24 hours. The latter cultures were used as the inocula for experiments. Cobamide producer isolates were pre-cultured by growing to saturation in medium lacking cobamide.

Growth measurements were performed in 200 µl cultures in 96-well plates (Corning #3598) and optical density at 600 nm (OD_600_) was recorded every 15 minutes in a Biotek Synergy 2 plate reader. Cobamide dose-response curves were acquired for six biological replicates by culturing in increasing concentrations of each cobamide with OD_600_ recorded every 15 minutes, as described previously (24). To determine maximum growth rates, the OD_600_ values were normalized to a 1 cm pathlength, blank subtracted in a replicate specific manner by subtracting the adjusted OD_600_ of the inoculum from the mean of the first two timepoints, and smoothed with 0^th^ order smoothing with 3 neighbors on either side in GraphPad Prism (v10.4.0). Values below 0.009 were eliminated and maximum growth rates were obtained using AMiGA (52). Dose response curves were plotted and fitted in GraphPad Prism.

To determine whether difference in dose response curves from the two dependent isolates were statistically significant, the fit parameters (EC50, Hillslope, fit top, and fit bottom) were compared via an ordinary two-way ANOVA followed by Sidak’s multiple comparisons test, with a single pooled variance using GraphPad Prism.

### Co-cultures of dependent isolates

Two independent experiments were performed on three independent cultures of each isolate, for a total of 6 biological replicates per condition. Each isolate was diluted to an OD_600_ of 0.0015 for monoculture and co-culture experiments. 1 ml cultures were grown in Masterblock (Greiner Bio-One) 96-well plates with 2 ml wells (covered with an AeraSeal (Excel Scientific) at 28°C with shaking at 800 rpm in a heated plate shaker (Southwest Science). 10 µl of culture were transferred into 990 µl of fresh medium containing the same cobamide conditions every 24 hours for 7 days. Serial dilutions and plating on R2A medium supplemented with 1 nM cyanocobalamin were done on days 0, 1, 3, 5, and 7 and colonies were counted after 48 hours of growth at 28°C. To determine if one isolate dominated the co-culture, we compared the mean *Ca*24A fraction at the final timepoint to a hypothetical value of 0.5 (representing equal proportions between the two species) via a one-sample t and Wilcoxon test in GraphPad Prism (v11.1.0).

### Producer supernatant assay

Four independent cultures of each producer were inoculated from glycerol stocks and grown to saturation in VL60 medium lacking cobamide. The cultures were sampled at exponential phase and stationary phase (16 and 36 hours for *Me*12 and 10 and 36 hours for *Pr*24B) and an aliquot was diluted and plated for viable cell counts (Fig. S1 C, D, E). The culture samples were centrifuged, and the supernatant was sterilized by filtration with a 0.22 µm syringe filter. Dependent pre-cultures were prepared as described above and inoculated into medium containing 10% v/v producer supernatant, a cobalamin solution, or water in a 96-well plate. The *Me*12 exponential phase supernatant was further diluted five-fold based on preliminary experiments that showed it contained saturating amounts of cobamide, and this dilution is accounted for in the number of dependent cells per producer cell ratio in Table 1. Serial dilutions and plating on R2A medium supplemented with 1 nM cyanocobalamin were performed at time 0 and after 24 hours of growth of each culture to determine the initial and final viable cell counts, respectively. Colonies were counted 48 hours after plating and the number of dependents supported by each producer was calculated by the following calculation: (Dependent CFU/ml gained when supplemented with supernatant – Dependent CFU/ml gained in no addition medium) / (Producer CFU/ml_final_ * dilution factor), where CFU/ml gained was calculated by subtracting the initial CFU/ml from the final CFU/ml.

To determine the impact of producer supernatant on growth of the dependents, data were compared with the no addition (NA) and cobalamin addition controls via ordinary one-way ANOVA followed by Šídák’s multiple comparisons test, with a single pooled variance on GraphPad Prism v10.5.0 (n=4 except in the case of *Ca*24A grown on *Me*12 exponential phase supernatant, where n=2, and *Ca*24A NA control, where n=3 due to sample loss, and in the case of *Ca*19 cobalamin control where n=3 due to a no growth outlier).

### Producer-dependent co-cultures

Dependents were pre-cultured as described above. Producer cultures (n=4) were inoculated from colonies and cultured in VL60 medium lacking cobamide for 24 or 48 hours for *Pr*24B and *Me*12, respectively. Producer cultures were washed by centrifugation and resuspension of cell pellets in fresh VL60 medium to remove spent medium containing cobamides. Both producer and dependent cultures were diluted to an OD_600_ of 0.1 in fresh VL60 medium and mixed 1:1 with another diluted culture or with fresh medium in quadruplicate. These mixtures were then inoculated 1:100 into 1 ml fresh VL60 medium in a 96-well plate with 2 ml wells. Cultures were serially diluted and plated to determine the viable cell count in the inocula and after 48 hours of growth at 28°C.

To determine the impact of growth in co-culture on each isolate, isolate growth was compared across conditions via an ordinary two-way ANOVA with main effects only, followed by Šídák’s multiple comparisons test, with a single pooled variance (n=4). To compare the effect of each producer on the growth of dependents in co-culture versus in the presence of producer supernatant, the number of dependent cells supported by each producer was determined by the following calculation: (Dependent CFU/ml gained in co-culture – Dependent CFU/ml gained in no addition medium) / Producer CFU/ml_final_, where CFU/ml gained was calculated by subtracting the initial CFU/ml from the final CFU/ml. To compare the ratios for each dependent isolate, we used an aligned rank transform (ART) ANOVA followed by Benjamini-Hochberg adjusted planned comparisons on R (v4.5.2).

### Tri-culture growth

Producers and dependents were prepared as described for the producer-dependent co-culture experiment. All cultures were adjusted to an OD_600_ of 0.1 and mixed in equal volumes to prepare the inoculum. 10 µl of inoculum was added to 990 µl of VL60 medium with no cobamide in a 96-well plate with 2 ml wells. Cultures were transferred and plated as described for the co-cultures of cobamide dependent isolates.

150 µl of culture were collected on days 1, 3, 5, and 7 and boiled to lyse the cells. This lysate was used to determine the total cobamide concentration via an *E. coli* bioassay that relies on growth of a cobamide-dependent strain (*E. coli* Δ*metE*) and a methionine-dependent control strain (*E. coli* Δ*metE* Δ*metH*), as described previously (24, 30).

### Genome sequencing and analysis

Genomic DNA was extracted from saturated cultures in VL60 medium using the Qiagen DNEasy Blood and Tissue kit adapted for Gram positive bacteria. Illumina libraries were prepared using 500 ng of input isolate DNA with the Illumina DNA prep kit with IDT® for Illumina® DNA/RNA index plates. Libraries were sequenced (2×150 paired end) on an Illumina NovaSeq X Plus instrument at Novogene (Novogene Corporation Inc., Sacramento, CA). Genomes were subsequently trimmed using Trimmomatic v0.39, assembled using SPAdes v3.15.3, and annotated using Prokka v1.14.5, and quality was assessed using QUAST v4.4 and CheckMv1.0.18 on the KBase platform as described previously (53, 54), and filtered for >95% completeness and <5% contamination. Amino acid sequences predicted by DRAM were used for downstream analyses (55). Genomes were taxonomically classified using GTDB-Tk (v2.3.2), and in the case of *Ca*19 and *Ca*24A, their whole genome sequences were compared using FastANI v0.1.3.

The genomes were functionally annotated using BlastKOALA (56) and cross-referenced with a list of HMMs for cobamide biosynthesis, cobamide-dependent functions, and independent alternatives to cobamide-dependent pathways (21, 25, 56). Transport genes and cobalamin-binding domains were left out of the analysis due to poor annotations, and rSAM-B_12_ proteins were omitted because this category included the Radical SAM superfamily, which includes but is not restricted to cobalamin-dependent rSAM enzymes. Additionally, genes that are not essential for cobamide biosynthesis or that are only found in archaea were not included.

Amino acid biosynthesis and small carbon source catabolism capability were assessed using GapMind (57). Pathway genes were considered to be present if they had at least a medium confidence hit as described previously (58). *Ca*19 was initially identified as a leucine auxotroph by GapMind due to one missing gene (*leuB*), but after manually searching the annotated genome, *leuB* was found in a different scaffold from other leucine biosynthesis genes, and *Ca*19 was reassessed as a leucine prototroph. Synthesis capacity of B vitamins other than cobalamin was assessed by evaluating the biosynthesis pathways with previously established thresholds (31, 32). Initially, BlastKOALA (56) annotations were searched for any instances of KEGG Orthology (KO) numbers from the previous work. After noticing incongruences between genomic predictions and experimental observations, Prokka annotations of the genomes were searched for KOs and corresponding Enzyme Commission (EC) numbers, and predictions for three of the four genomes were reassessed after additional biosynthesis genes were detected.

### Experimental validation of B vitamin production and dependence

Triplicate pre-cultures of each isolate were inoculated from colonies into VL60 medium containing 1 nM cobalamin and grown to saturation. Pre-cultures were passaged 1:100 into 1 mL VL60 medium lacking one of biotin, niacin, thiamin, or riboflavin, or lacking all vitamins, or with all vitamins as a positive control. Passaged cultures were grown in 96-well plates with 2 mL wells for 24 hours (*Ca*19, *Ca*24, *Pr*24B) or 48 hours (*Me*12), after which growth was evaluated with OD_600_ measurements. Passaging (1:100) was done into fresh medium continued for each condition until no growth (OD_600_ < 0.009) was observed, or for up to four passages, at which vitamins from the pre-cultures would have been diluted to sub-picomolar concentrations.

## Data Availability

The sequencing data generated during the current study are available in the NCBI GenBank repository under Biosample Accession IDs SAMN51232832 – SAMN51232835. Code for genome analysis based on enzyme commission numbers is available on GitHub (https://github.com/stqngu/genome_annotation_processing).

## Acknowledgements

We are grateful to Will Ludington for his advice on studying competition at different cobamide concentrations and for pointing us to key literature, Christine Qabar for assistance with experimental troubleshooting and statistical analyses, Hans Carlson for assistance with sequencing efforts, Eleanor Wang for assistance with genome analysis, Alexa Nicolas for compiling genome analysis resources, and Zachary Hallberg for statistical recommendations. We thank Ashley Eng, Cami Scantlin, Dennis Suazo, Eleanor Wang, and Zachary Hallberg for critical reading of the manuscript.

This work was funded by the National Institutes of Health (NIH) Grant R35GM139633 (M.E.T.). H.I.A. was supported by the Bay Area NSF RaMP (Grant # 2216550).

