## Supplemental Figures 1-5 for "Cobamide-based interactions between soil bacteria can be predicted based on monoculture growth"

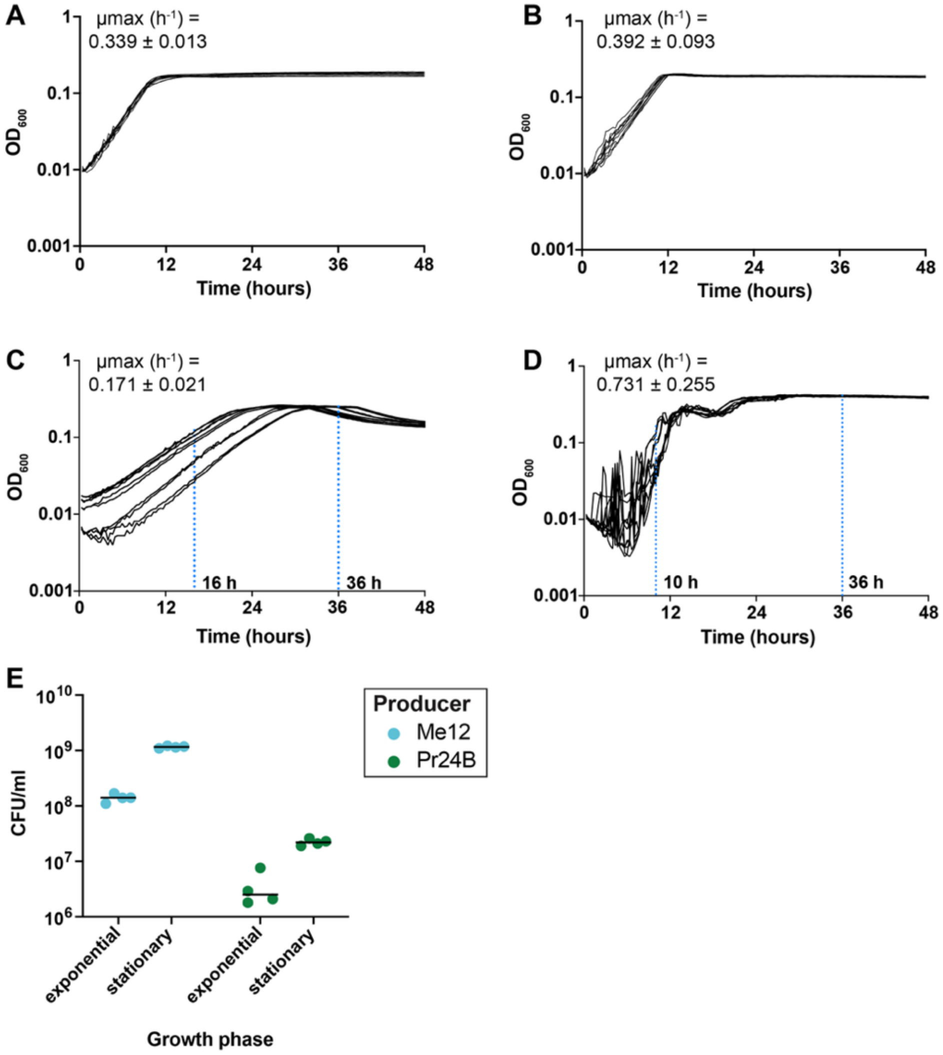


**Figure S1. Growth curves and viable cell numbers of isolates in this study.** Growth curves of (A) *Ca*19 and (B) *Ca*24A in VL60 medium with 1 nM cobalamin, and (C) *Me*12 and (D) *Pr*24B in VL60 medium without added cobalamin. The $OD_{600}$ readings were blanked and pathlength corrected as described in the methods. Lines show individual measurements of 5 biological replicates, each with 2 technical replicates. Dotted vertical blue lines show the supernatant sampling timepoints during the exponential and stationary phases of growth. The growth rate for each isolate was determined using AmiGA v.3.0.4 on the blanked, pathlength corrected, and smoothed growth curves and the mean maximum growth rate (μmax) and standard deviation (n=10) for each isolate are shown above each growth curve. (E) Producer CFU/ml at the supernatant sampling timepoints for exponential and stationary phase.

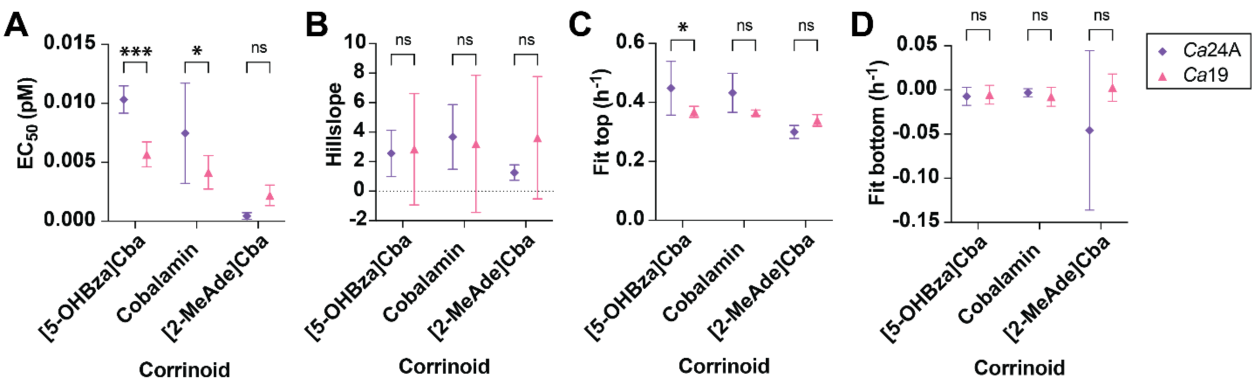


**Figure S2. Comparison of least squares fit parameters for dose response curves**. Parameters for dependent isolate dose response curves shown in Figure 2A, D, G. (A) Cobamide concentration that leads to half maximal growth rate ($EC_{50}$), (B) hillslope, (C) fit bottom, and (D) fit top for each isolate on [5-OHBza]Cba, cobalamin, and [2-MeAde]Cba highlight the differences in their cobamide preferences. Points and error bars show the mean and standard deviation for each value (n=5 or n=6). Fit parameter data were generated and compared on GraphPad Prism v10.5.0 via an ordinary two-way ANOVA followed by Sidak's multiple comparisons test, with a single pooled variance. One asterisk represents $p<0.05$, two asterisks $p<0.01$, three asterisks $p<0.001$, and four asterisks $p<0.0001$, while 'ns' is used for non-significant ($p>0.05$).

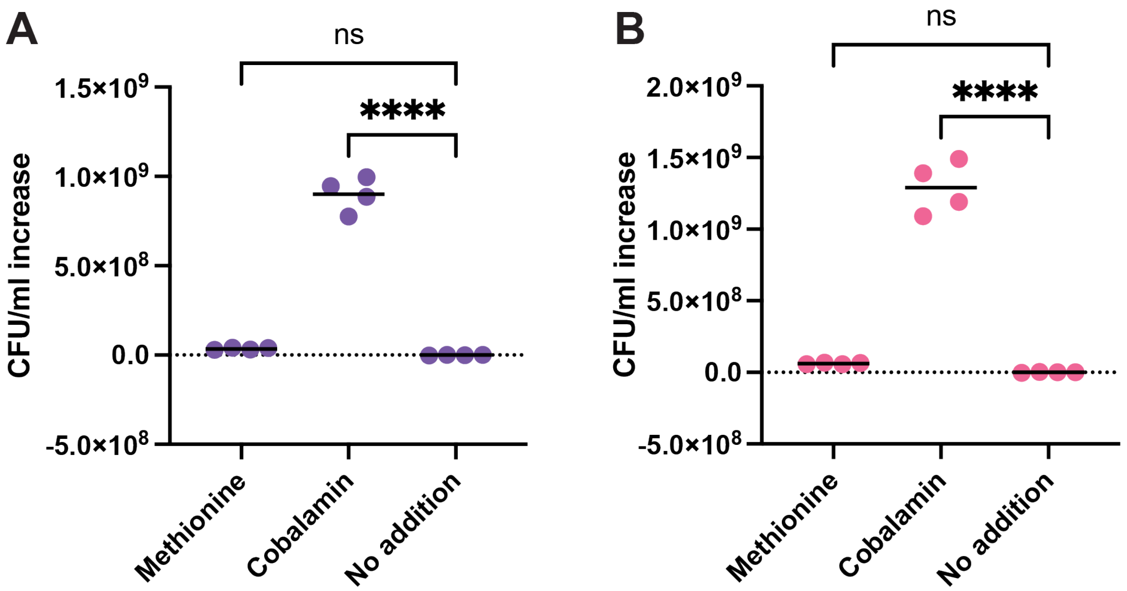


**Figure S3. Growth of dependent isolates (A) *Ca*24A, (B) *Ca*19 on 0.15 mg/ml methionine, 1 nM cobalamin, and no addition control.** Points show individual replicates and horizontal lines show the mean CFU/ml increase for each isolate in each condition. Data points were compared via ordinary one-way ANOVA followed by Dunnett's multiple comparisons test, with a single pooled variance. One asterisk represents $p<0.05$, two asterisks $p<0.01$, three asterisks $p<0.001$, and four asterisks $p<0.0001$, while 'ns' is used for non-significant results.

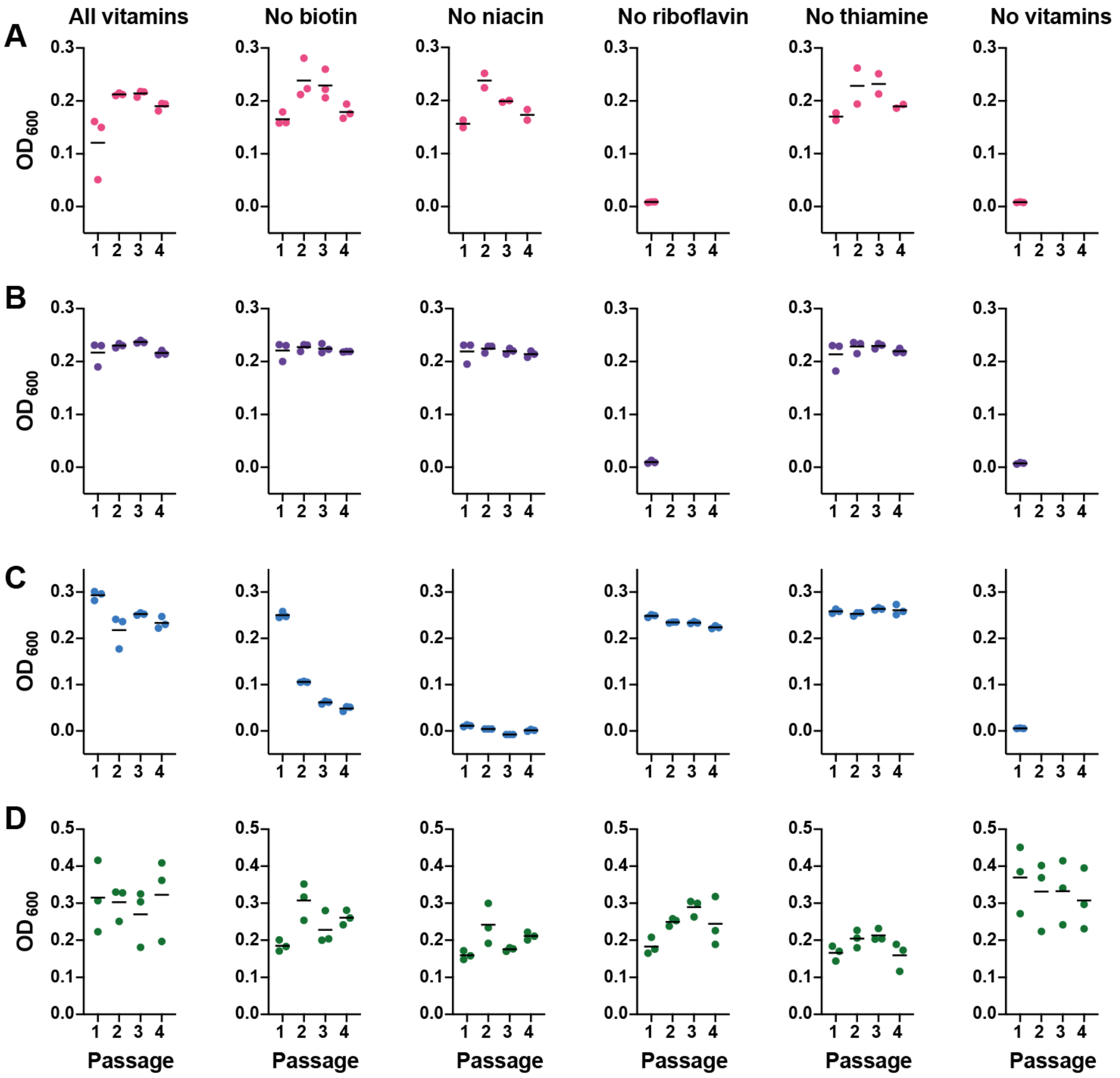


**Figure S4. Experimental confirmation of vitamin production and dependence for biotin, niacin, riboflavin, and thiamine.** OD600 after 24 hours of growth (*Ca*19, *Ca*24, *Pr*24B) or 48 hours of growth (*Me*12) is shown for (A) *Ca*19, (B) *Ca*24A, (C) *Me*12, and (D) *Pr*24B**.** Biological replicate cultures (n=3, one no-growth outlier was removed in *Ca*19-niacin and Ca19-thiamine) were precultured in VL60 medium containing all vitamins and subsequently passaged four times into VL60 medium lacking one of biotin, niacin, riboflavin, or thiamine. Only one passage is shown for instances where the isolate stopped growing immediately after the first passage.

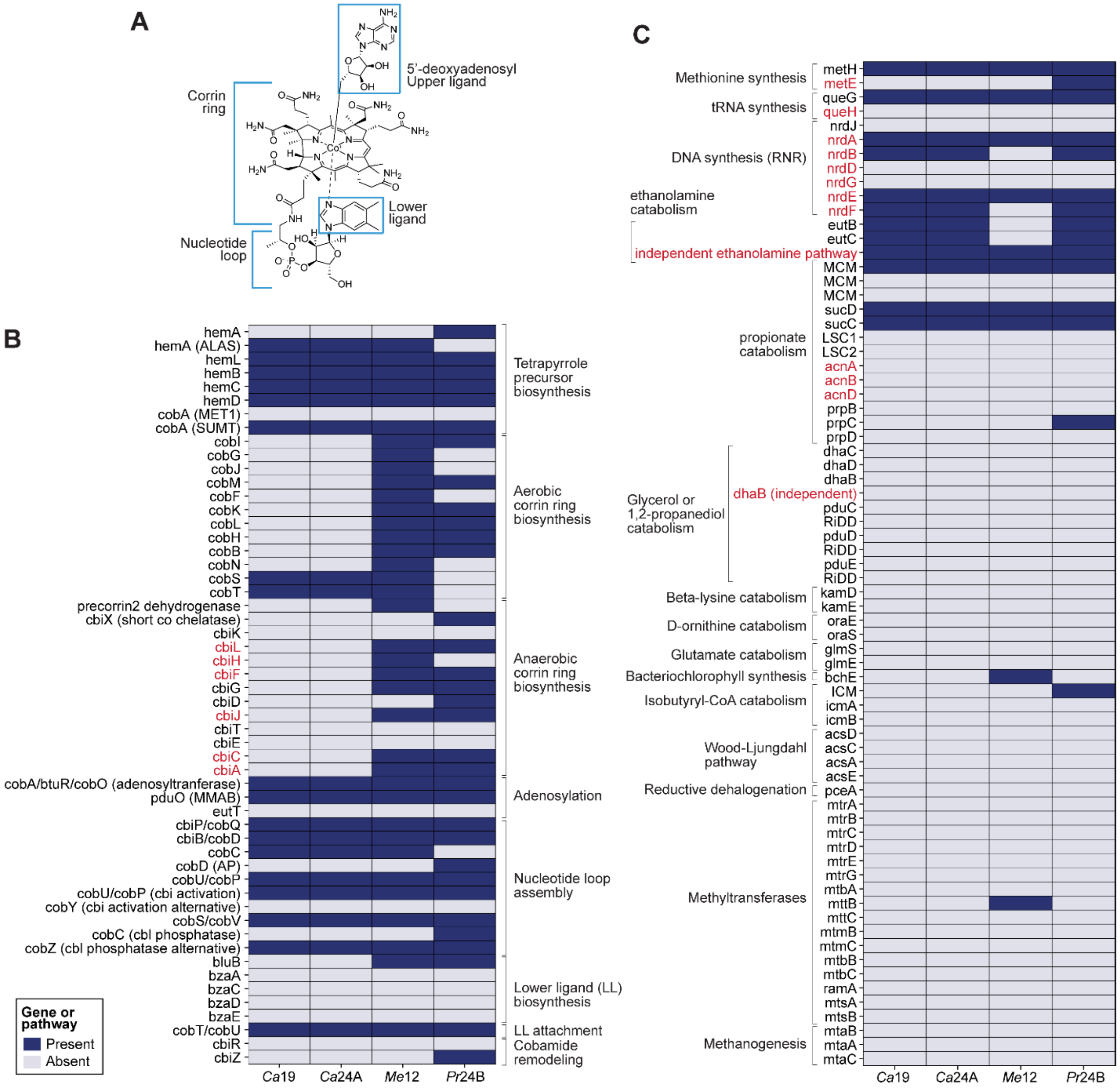


**Figure S5. Presence or absence of corrinoid biosynthesis and corrinoid-dependent genes and pathways.** (A) Structure of adenosylcobalamin ($B_{12}$) with major components labeled. (B) Presence and absence of corrinoid biosynthesis genes listed on the left are indicated by the colored boxes. Gene names in red function in anaerobic corrin ring biosynthesis that share KOfams with genes in aerobic corrin ring biosynthesis. (C) Corrinoid-dependent genes or pathways are shown in black text and corrinoid-independent alternatives are shown in red text. Genes are grouped by the type of metabolism in which they are involved. For more details regarding the genes considered here, see Table S2.
